# Temperate grasslands facing heatwaves: species diversity buffers effects on shoot growth but not on leaf parameters

**DOI:** 10.1101/2025.11.04.686479

**Authors:** Andreu Cera, Sophie Brunel-Muguet, Servane Lemauviel-Lavenant

## Abstract

Current and future heatwaves threaten temperate grasslands. Plant temperature regulation, photosynthesis and therefore production are exposed. As with drought, species diversity could buffer the negative effects of heatwaves. Our main objective was to test the effect of diversity on grassland responses to heatwaves by comparing mixtures with monocultures. We conducted a greenhouse experiment to simulate four thermo-protocols: i) control ii) two mild heatwaves, iii) a severe heatwave, and iv) recurrent scenario with mild and severe heatwaves. We analyzed production at mowing time, resistance and recovery of species calculated from shoot growth, and foliar parameters sensitive to heatwaves such as leaf temperature and maximum quantum efficiency of Photosystem II. A severe heatwave induced negative effects on production and leaf parameters, whereas mild heatwaves not. Recurrent heatwaves induced negative cumulative effects on production and acclimation of Photosystem II. In this context, mixtures were more productive than monocultures average, as overyielding; certain species showed higher resistance or recovery in mixtures than in monocultures; but there was no diversity effect on leaf parameters. Our results suggest that leaf parameters alone cannot reflect the resilience of plants exposed to heat and that plant diversity can give temperate grasslands a higher capacity to cope with heatwaves.

## Introduction

Extreme weather events are threats for plants and addressing them is essential for the management of species and ecosystems (Maxwell *et al*., 2019). Heatwaves, one of such extreme weather events, are defined by the occurrence of abnormally high temperatures for several days (Perkins & Alexander, 2013). There is compelling evidence that heatwaves have become longer, more frequent and intense in most land regions since the 1950s (IPCC 2023, 6^th^ Assessment Report). However, the impact of heatwaves on plants has been under-researched compared to other extreme weather events such as droughts (Breshears *et al*., 2021).

Heatwaves affect biomass production in temperate grasslands (Langworthy et al., 2020; Wang et al., 2008). However, the impact varies depending on the timing of their occurrence. In temperate-climate regions of North hemisphere, heatwaves can occur from April to September (Lavaysse et al., 2019), thus affecting a wide window within the phenological stage of plants and the state of ecosystems, particularly the availability of water in the soil. Spring or autumn heatwaves have rather neutral effects on production of temperate grassland species (De Boeck et al., 2011), whereas summer heatwaves cause more severe damage (Wang et al., 2008). This seasonal effect is not only related to the margin between plant thermotolerance and the temperatures reached during heatwaves (Kitudom et al., 2022), but also and mainly, according to some other authors, to soil moisture conditions (De Boeck et al., 2011; Dreesen et al., 2012; Reynaert et al., 2025). Water availability is a key factor in coping with heatwaves, as plants respond to heat episodes by increasing transpiration to lower internal temperatures (Körner, 2013). However, it is still unclear whether plant thermotolerance or soil water availability is the main driver of the negative effects of heatwaves on production of temperate grassland species. When heatwaves coincide with droughts, water shortage is the main stress for temperate grassland species (De Boeck et al., 2011). This type of extreme weather events (combined drought and heatwave event) reflects an increasing trend due to global change. However, only up to 20% of heatwaves coincide with droughts (Mukherjee & Mishra, 2021). The negative impact of combined drought and heatwave events on production of temperate grassland species is clear (Milbau et al., 2005; Poirier et al., 2012; Volaire et al., 2020), especially in dry temperate regions where soil moisture is more frequently limited (e.g. Southern European regions). Nevertheless, it seems important to disentangle the effect of heatwaves alone on temperate grasslands in non-water-limited systems and to resolve the controversy about the main cause of induced negative effects.

Heat events can recur in the same year in different seasons or even in the same season (Christidis et al., 2015). This leads to a need to better consider the recurrence of heat events in order to understand the heatwaves impact on grasslands. Recurrent events can produce positive or negative effects on the performance of plants, which have developed the ability to modify their responses to an event if previously exposed to an event of the same or different nature through memory processes (Ogle et al., 2015). Faster, more intense and/or more sensitive responses to events following prior exposure(s), define the so-called “priming effects” (Bäurle, 2016), which to some extent allow for event acclimation (Smith & Dukes, 2013). Such positive effects, through priming-related processes, can occur in grassland species for example with the development of new thermotolerant leaves (Davies et al., 2018) or a plasticity of photosynthesis tolerance (French et al., 2019; Zhu et al., 2018). Conversely, cumulative negative memory effects have also been reported in grassland species following recurrent heat events (Lemoine et al., 2018; Notarnicola et al., 2021; Vandegeer et al., 2020). Therefore, while the effects of heatwaves on grasslands are well characterized at the species and community level, their recurrence may alter these responses, making this feature key for representing heatwaves-induced effects in experimental studies.

Multi-species assemblages can achieve overyielding, where the mixture performance exceeds the average of monocultures, but also transgressive overyielding, where the mixture performance exceeds that of the best performing monoculture (Hector et al. 2002). Higher levels of diversity in plant communities have been associated with better performance and tolerance to extreme weather events, such as drought (Grange et al., 2021; Haughey et al., 2023; Hofer et al., 2016), when compared to monocultures, even in the context of recurrent events (Chen et al., 2022). The overyielding observed in drought experiment, is acquired through synergistic interactions between species, such as the facilitation through legumes atmospheric N fixation or complementarity (Haughey et al., 2023), or through the sampling effect (Loreau, 1998), which means the outperformance of certain species during or after the extreme weather event (Hofer et al., 2016). These results provide evidence that high levels of biodiversity can buffer ecosystem functioning under extreme weather events, as described by the biological insurance theory (Loreau et al., 2021). Apart from evidence from drought research, how plant species diversity can modulate the effect on grasslands in the context of heatwaves remains unexplored. In particular, plant responses such as resistance (i.e., the maintenance of shoot growth during the event) and recovery (i.e., the ability to recuperate shoot growth after the event) or traits that explain plant performance under heatwave-induced effects, such as maximum quantum efficiency of Photosystem II (Geange et al., 2021) or foliar temperature (Blonder & Michaletz, 2018) remain to be investigated. A positive effect of plant diversity but also of specific plant traits that ensure heat buffering, could contribute to the development of climate-resistant or resilient grasslands (Reynaert et al., 2025).

Our aim was to test the effect of species diversity on the response of grasslands to heatwaves by comparing plant mixtures with monocultures. To this end, we conducted a greenhouse experiment that simulated the effects of recurrent heatwaves (HW) - two mild HW and one severe HW - on sown, mowed grasslands under non-nutrient and water-limiting conditions (Figure 1). Four temperate grassland species, *Festuca rubra, Lolium perenne, Lotus corniculatus* and *Plantago lanceolata*, were selected because of their potentially contrasting responses to HW in terms of tolerance to extreme weather events (Grime *et al*., 2014) and because they belong to the main functional groups of grasslands (grasses, legumes and forbs). We then analyzed the effect of plant diversity by comparing the performance of monocultures with that of grass-legume and multi-species mixtures throughout the experiment, and especially after mild and severe HWs. We analyzed aboveground production related to mowing, but also leaf parameters, sensitive to heatwaves such as foliar temperature, and maximum quantum efficiency of Photosystem II, to assess species resistance and recovery. We formulated a hypothesis based on the biological insurance theory (Loreau *et al*., 2021), assuming that grassland species show contrasting responses to heatwaves and, consequently, that assemblages of different species can buffer heatwaves-induced impairments compared to monocultures of the same species. Thereby, multi-species mixtures would be more productive than monocultures, plants growing in mixtures would be more resistant and/or have higher recover to heatwaves than growing in monocultures.

**Figure 1.**
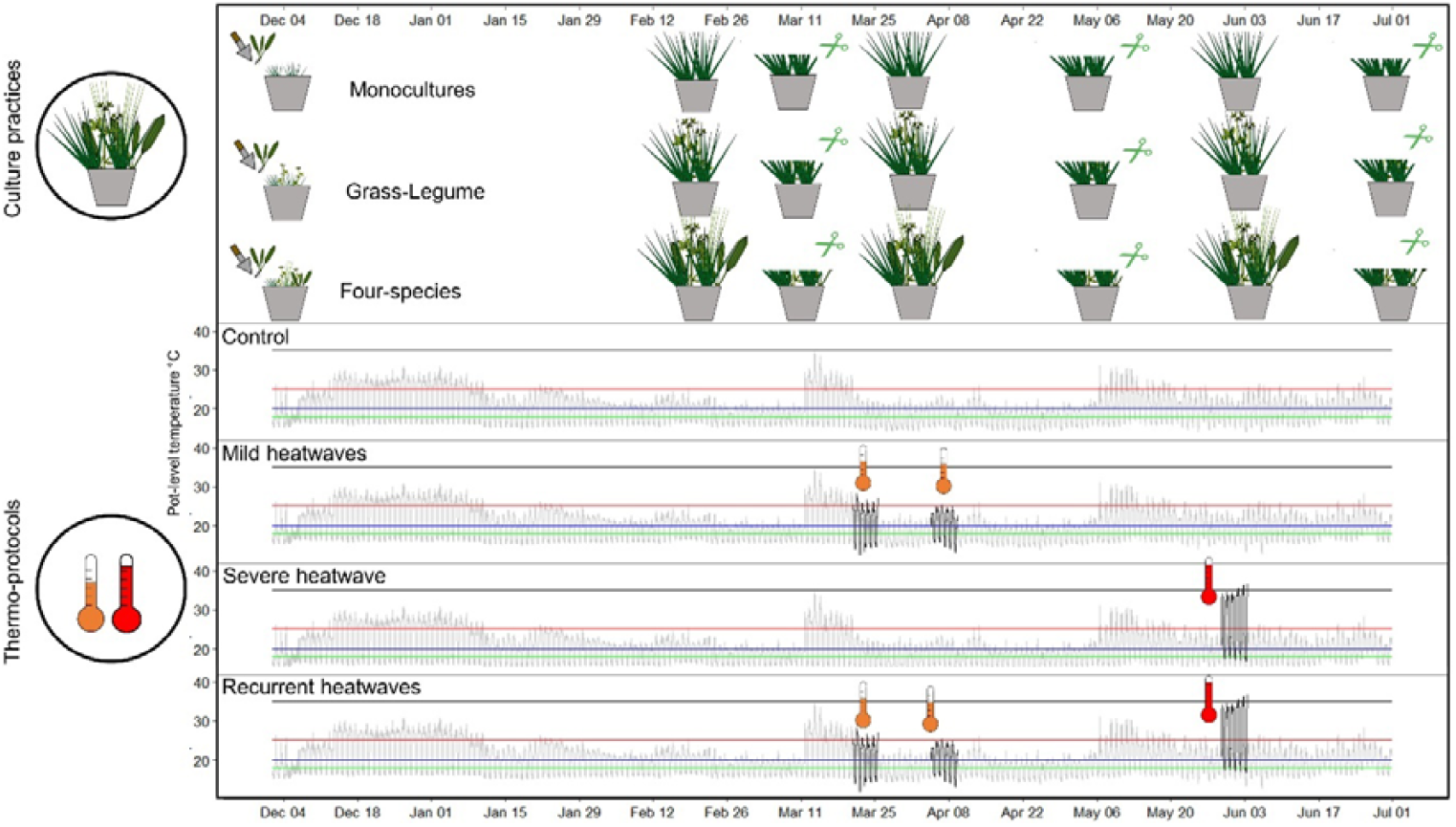
Experimental design and observed air temperatures. Cultures practices, which involve three type of cultures (monocultures, grass-legume and four-species mixtures), are plant transplantation on December 4, and three mowing on March 11, May 6 and July 1. Thermo-protocols consist of the application or not of two types of heatwaves, namely mild heatwaves on March 25 and April 6, and severe heatwave on June 3. Temperatures were measured at pot level. Control unit temperatures are grey lines and heatwave unit temperatures are black lines.

## Materials and Methods

### 2.1 Studied species and selected varieties

The species used in this study are temperate grasslands species commonly sown in Europe i.e. *Lolium perenne* L., *Festuca rubra* L., *Lotus corniculatus* L., and *Plantago lanceolata* L.. The varieties used in the greenhouse experiment, were provided by two seed companies, “Elixir” for *L. perenne* was supplied by Cerience, and “Bardance” for *F. rubra*, “Lotar” for *L. corniculatus* and “Captain*”* for *P. lanceolata* were supplied by Barenbrug.

### 2.2 Pots preparation and location

The experiment was carried in a greenhouse at Université de Caen-Normandie in France (49°11′09 N, 0°21′32 W). Seeds of each studied species were germinated separately in rectangular plastic pots filled with three-part of osmotic water and one-part of perlite on November 24^th^, 2023, in the greenhouse at 20°C (16 hours daylight with artificial light) and 16°C (8 hours night). Pots were covered with aluminum foil until the emergence of cotyledons, and then uncovered progressively according to the emergence of each species. On December 4^th^, 2023, seedlings were transplanted to cylindric pots of 4 L with quarry silica sand (Silica BB 0.8-1.8 DS:BL, Sibelco, Mios, France). For each pot, we transferred 2 to 3 individuals into each of the 8 holes provided to install 8 plants per pot. Fifteen days after transplantation, pots were thinned to one plant per hole. All plants were kept well-watered throughout the experiment with a Hoagland solution ¼: 1.25 mM Ca (NO_3_)_2_ 4 H_2_O, 1.25 mM KNO_3_, 0.25 mM KH_2_PO_4_, 0.5 mM MgSO_4_, 0.2 EDTA (NaFe 3 H_2_O), 0.014 mM H3B3, 0.005 mM MnSO_4_ H_2_O, 0.003 mM ZnSO_4_ 7H_2_O, 0.0007 mM CuSO_4_ 5H_2_O, 0.0007 (NH_4_)_6_ Mo_7_O_24_, 0.0001 CoCl_2_. The nutrient solution was administrated with an automatic closed circulation system. The solution was renewed or added every 3 days and each pot received a total of 300 mL per day, distributed in 5 times. Pots were installed on mobile rack that were moved once time per month to avoid microhabitat effects.

### 2.3 Experimental design

The experiment consisted in simulating heatwave scenarios and grassland management. We prepared three types of grassland cultures to study the effect of species diversity on grassland performance: monocultures for each of the four species; a grass-legume mixture of *Lolium perenne* and *Lotus corniculatus*; and a mixture of the four species. All plants were grown into the greenhouse from germination until final harvest and subjected to a standard thermoperiod (20/16°C, d/n) (Figure 1). To simulate heatwaves, we designed four thermo-protocols, based on those previously designed and tested in crops species in similar semi controlled conditions (Magno Massuia de Almeida *et al*., 2023). On March 11, 2024, 16 weeks after transplantation of plants to sand pots, we divided the number of pots to four thermo-protocols: i) Control conditions i.e. standard thermoperiod; ii) Mild heatwaves i.e. the cultures were subjected to two events of mild warning (25/18°C, d/n) for five days on the 16^th^ week and on the 18^th^ week after transplantation; iii) Severe heatwave i.e. the cultures were subjected to a single event of severe warning (41.5/19.5°C d/n) for five days from the 26^th^ week after transplantation; iv) Recurrent heatwaves i.e. the cultures were subjected to the three heatwaves (two mild and one severe heatwaves). Heatwaves were applied in a separate unit in the greenhouse. Temperature within the control and heat units were controlled by the air-conditioning system (Kairos, Anjou Automation, Mortagne sur Sèvre, France). We controlled chamber temperature at the pot-level with a temperature sensor (Aranet4, SAF Tehnika JSC, Riga, Latvia). Plants subjected to heatwaves received the same amount of water as plants not subjected to heatwaves. The pot mass loss was recorded to each heatwave to infer the induced water shortage. We applied three mowing at 5 cm to all pots at three dates: March 11^th^, 2024; May 6^th^, 2024; and last mowing and harvest on July 1^st^, 2024 to simulate mowing in a Normandy context. For each thermo-protocol, we had four replicates per monoculture, six replicates of grass-legume mixture and eight replicates of four-species-mixture, making a total of 120 pots. Pests, mainly aphids and thrips, were controlled by spraying or washing the leaves with tap water.

### 2.4 Shoot growth, yield and leaf traits

Maximum shoot length (SL) was measured, from the base of plant to the most distant leaf, monthly and before and after a heatwave using a metallic millimeter straightedge. Shoot growth (SG) between two times (days) t1 and t2 was calculated as:

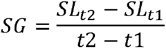

Aboveground mass at 5 cm was assessed for each pot after each mowing. At the final harvest (July 1), total aboveground mass per species was assessed for each pot. Biomass was dried to a constant mass at 50°C and weighed in a precision scale (3200 g/0.01 g, MS3002S/01, Mettler Toledo, Columbus-OH, USA).

Leaf temperature was measured as it provides information on how plants manage the excess energy due to heatwaves through dissipation (Körner, 2013). Leaf temperature was measured in the upper third of the canopy. We selected leaves in good condition and without any wounds. The measurement was conducted at one point on the leaf with a thermal imaging camera (WB-80, Voltcraft, Hirschau, Germany) on the fifth day of heatwaves events between 10-12 am. We measured one leaf for each species per pot both for pots subjected to the mild and severe heatwaves and the control pots.

The maximum quantum efficiency of Photosystem II, widely considered as a sensitive indicator of plant photosynthetic performance (Geange *et al*., 2021), was measured in the upper third of the canopy, on good conditions leaves without any wounds. Fv/Fm is presented as a ratio of variable fluorescence (Fv) over the maximum fluorescence value (Fm) and is calculated as (Fm - Fo)/Fv. This ratio varies between 0 and 1. Lower values are usually observed under stressful conditions. The measurement was conducted at one point on the leaf using a leafclip with an excitation fluorimeter (PocketPea, Hansatech Instruments, Norfolk, United Kingdom). The measured points were 15 minutes in the dark before excitation with 2800 µmol m-2 s-1 intensity. We measured Fv/Fm on the fifth day of heatwave events between 10-12 pm on one leaf of each species per pot for both pots subjected to the mild and severe heatwaves and the control pots.

### 2.6 Data analyses

To assess pot mass loss and evaluate water shortage per each heatwave, we carried out ANOVAs per using linear models with chamber (heatwave or control) as categorical fixed factor. We run linear models using glmmTMB function in glmmTMB package (Brooks *et al*., 2024).

To identify the determinant factors in yield, we used cumulative yield through the experiment, 1^st^ mowing yield, 2^nd^ mowing yield, 3^rd^ mowing yield and indices of overyielding and transgressive overyielding. Cumulative yield is the sum of the yield above 5 cm through the three mowing. The overyielding and transgressive overyielding indices were calculated to evaluate whether higher diversity allows a better production than the mean of the monoculture production or than the most productive monoculture in our experiment (see Loreau 1998). Overyielding index (*D*_*mean*_) and transgressive overyielding (*D*_*max*_) index were calculated from cumulative yield and each mowing yield as:

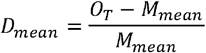

where *O*_*T*_ is the observed mass of a given mixture, and *M*_*mean*_ is the mean mass produced by monocultures of any of the species in this mixture. A *D*_*mean*_ *> 0* indicates an overyielding.

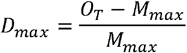

where *O*_*T*_ is again the observed mass of a given mixture, and *M*_*max*_ is the mass produced by the most productive monoculture of any of the species in this mixture. A *D*_*max*_ *> 0* indicates a transgressive overyielding (Loreau, 1998).

To evaluate the impact of mild and severe heatwaves, we used indices of resistance and recovery to evaluate the response on shoot growth of species in our experiment. Shoot growth was calculated two times: i) during heatwave events, calculated between the day before the heatwave and the last day of the heatwave; and ii) after heatwave events, calculated between the last day of the heatwave and the day before the next harvest. We calculated Z-score relative to control conditions using shoot growth during heatwave as resistance index, and used shoot growth after heatwave as recovery index.

To assess changes in yield, we carried out ANOVAs using linear models with mild heatwaves, severe heatwave, culture (*F. rubra* monoculture, *L. perenne* monoculture, *L. corniculatus* monoculture, *P. lanceolata* monoculture, grass-legume mixture, four-species-mixture) and the interactions among them as categorical fixed factors. We added previous mowing yield as a covariable to evaluate the 2^nd^ mowing yield and 3^rd^ mowing yield, to take into account the variability of each pot. We run linear models using glmmTMB function in glmmTMB package (Brooks *et al*., 2024). When differences were statistically significant, multiple comparisons between levels of fixed factors were assessed using emmeans function in emmeans package version 1.4-13 (Lenth *et al*., 2023) and multcomp function (Hothorn *et al*., 2025). To conclude on overyielding and transgressive overyielding from the indices, we carried out a contrast against 0 to evaluate whether *D*_*max*_ *>0* or *D*_*mean*_ *>0* with µ=0 using emmeans function for each treatment (combination of thermo-protocol and type of culture). To evaluate resistance and recovery, we carried out a contrast against 0 to evaluate whether *resistance*≥*0* or *recovery* ≥*0* with µ=0 using emmeans function for each treatment. We also carried out ANOVA using linear models with type of heatwave, culture (all possible levels) as categorical fixed factors for each species. To evaluate the responses of leaf parameters to mild heatwaves and severe heatwave, we carried out ANOVA using linear models with type of heatwave, culture (all possible levels) as categorical fixed factors for each species.

All calculations, statistical analyses and graphics were carried on using R v.4.4.2 (R Core Team, 2023).

## Results

### Pot-level temperature and pot mass variation

The first mild heatwave reached a maximum of 28.15 °C and a mean of 24.13 °C during daylight hours at the pot-level (Figure 1) while during the same period, the temperature in the control unit reached a maximum of 24 °C and a mean of 21.48 °C. There was not significant mass loss on pots from both units indicating an absence of heatwave-induced drought (Figure S1, Table S1). The second mild heatwave reached a maximum of 25.35 °C and a mean of 23.00 °C during daylight hours at the pot-level (Figure 1) while during the same period, the temperature in the control unit reached a maximum of 23.55 °C and a mean of 21.47 °C. There was not significant mass loss on pots from both units (Figure S1, Table S1). Eventually, the severe heatwave reached a maximum of 36.6 °C and a mean of 30.68 °C during daylight hours at pot-level (Figure 1) while during the same period, the temperature in the control unit reached a maximum of 24.15 °C and a mean of 21.52 °C. A significant mass loss in pots from the heat unit was observed, with an average reduction of 7% compared to the ones in the control unit, indicated a water loss (Figure S1, Table S1), without differences between thermo-protocols.

### Effects of thermo-protocols and types of cultures on yield

The two mild heatwaves (i.e., mild HW thermo-protocol) and the severe one (i.e., severe HW thermo-protocol) appeared to have no significant negative impact on total yield (cumulating all mowing yields), while when combined (i.e., recurrent heatwaves thermo-protocol) the effects were significant negative (Figure 2, see ANOVA summary on Table S2). The recurrent heatwaves thermo-protocol was the least productive thermo-protocol with similar production to the ones under control in contrast to mild heatwaves or a severe heatwave that both led to similar higher yields at the end of the experiment. Regarding the effect of heatwaves at 2^nd^ and 3^rd^ mowing yield, mild heatwaves had no impact on yield at 2^nd^ mowing (Figure S2), whereas yield at 3^rd^ mowing was significantly impacted. The recurrent heatwaves were the least productive thermo-protocol at 3^rd^ mowing as they reduced the yield 1.21-fold lower than the severe heatwave alone, which is the second least productive. Mild heatwaves and control conditions had similar effect on production at 3^rd^ mowing, and induced higher production than severe heatwave and recurrent heatwaves thermo-protocols Culture type was determinant on yield at each mowing and for the cumulative yield (P<0.05, Figure 2, see ANOVA summary on Table S2). Whatever the thermo-protocol, the most productive cultures (cumulative yield) were the grass-legume mixture, the monoculture of *L. perenne* and the four-species mixture (Figure 2). The least productive culture was the monoculture of *L. corniculatus*, 1.55-fold lower than grass-legume culture, and the monoculture of F. rubra, 1.74-fold lower than grass-legume mixture. Monoculture of *P. lanceolata* was more productive than monocultures of *F. rubra* and *L. corniculatus*, similar to monoculture of *L. perenne* and the four-species mixture, but 1.14-fold lower than grass – legume mixture. The yield differences among culture types were not stable through the experiment. At the 1^st^ mowing (Figure S2), yields among the culture types were similar except for the monoculture of *F*.*rubra* which displayed a lower value. At the 2^nd^ mowing (Figure S2), the monoculture of *L. perenne* was the most productive, followed by the grass-legume mixture, the monoculture of *F*.*rubra*, the monoculture of *P*.*lanceolata* jointly with the four-species mixture, and the monoculture the *L*.*corniculatus* being least productive. At the 3^rd^ mowing (Figure 2), the most productive were the two mixtures and the monoculture of *L*.*perenn*e, followed by the monoculture of *P*.*lanceolata*, the monoculture of *F*.*rubra* and the monoculture of *L*.*corniculatus* being the least productive on average of the thermo-protocols.

**Figure 2.**
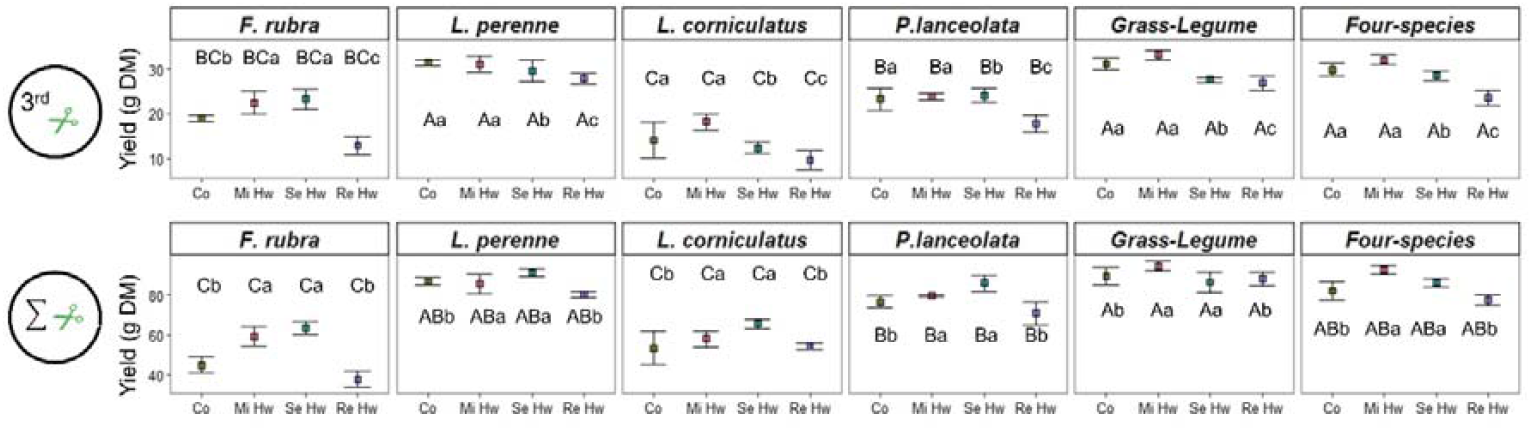
Yield (g Dry Mass) for 3^rd^ mowing and sum of all mowing, culture type (monocultures, grass-legume and four species mixtures) and thermo-protocol (Co: control, Mi Hw: mild heatwaves, Se Hw: severe heatwave, Re Hw: recurrent heatwaves). Means and standard error are shown. Capital letters indicate significant difference among culture types after multiple comparisons, and minor letters indicate difference among thermo-protocols for each culture type (p<0.05).

### Overyielding through the experiment

Overyielding (D_mean_ > 0) was evidenced in our experiment for all the thermo-protocols (Figure 3A, see table S3 for t-test summaries), except for the grass-legume culture subjected to severe heatwave at 3^rd^ mowing. Nevertheless, transgressive overyielding (D_max_ > 0) has not been evidenced in our experiment (Figure 3A, see table S4 for t-test summaries) except for the grass-legume culture subjected to mild heatwaves at 2^nd^ mowing (*P*<0.05).

**Figure 3.**
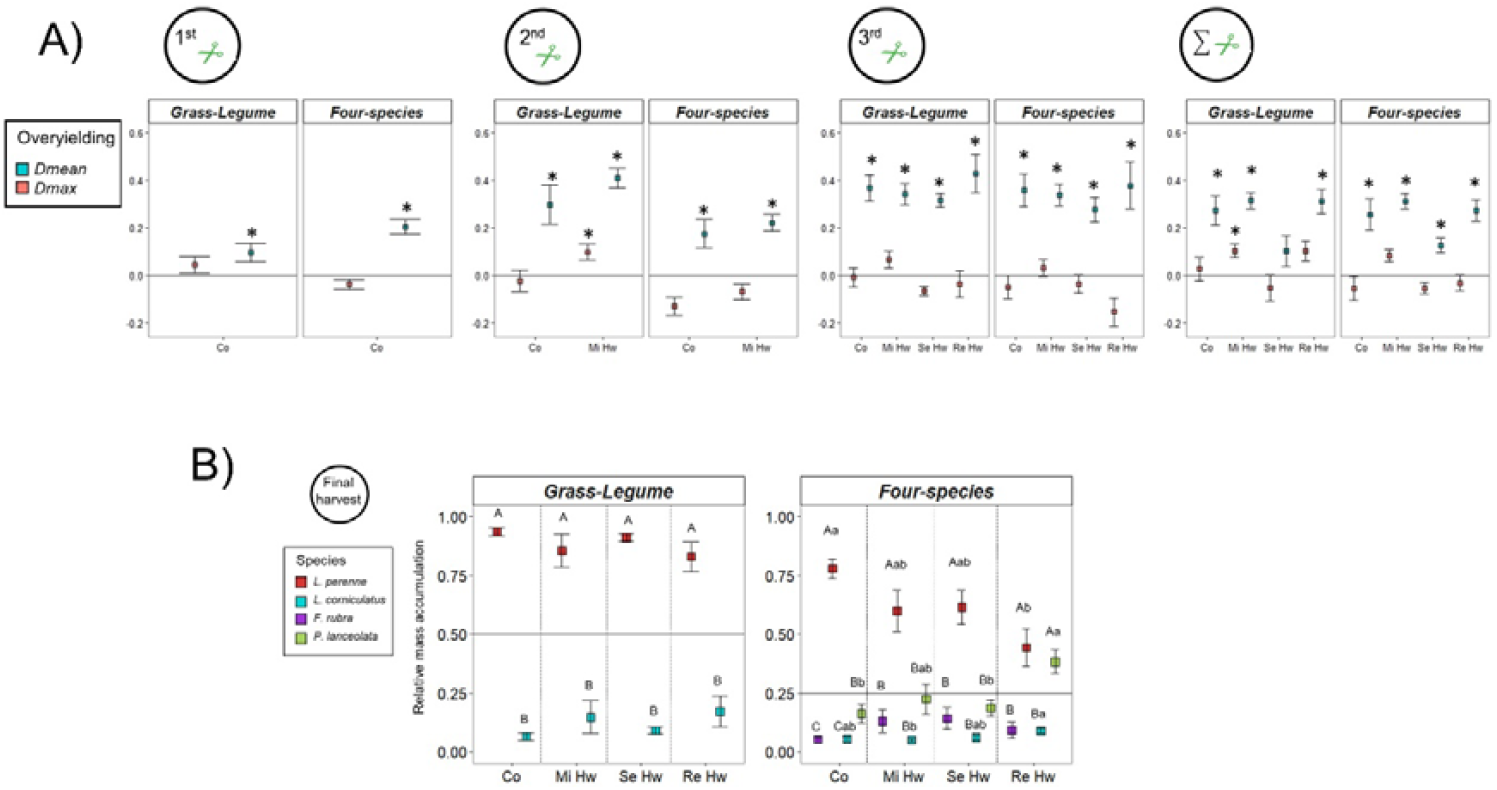
Overyielding analysis through the indices of overyielding and transgressive overyielding that compare mixtures with monocultures. Thermo-protocols are **Co**ntrol (Co), **Mild** heatwaves (Mi Hw), **Severe** heatwave (Se Hw), **Re**current heatwaves (Re Hw). On the section A, overyielding (*D*_*mean*_) and transgressive overyielding (*D*_*max*_) indices at 1^st^ mowing, 2^nd^ mowing, 3^rd^ mowing and for the total (sum of all mowing) analyzed for the grass-legume mixture and the four species mixture. Asterisk indicate whether Dmean and Dmax are larger than zero based on t-tests. On the section B, Relative mass of each species in mixtures (Grass-Legume and Four-species) calculated by using the aboveground mass at the final harvest. From multiple comparisons (P<0.05), capital letters are significant differences within thermo-protocol among species for each type of mixture, and minor letters are significant differences within mixtures among thermo-protocols for each species.

Overyielding in the grass-legume mixture was mainly due to the overperformance of *L. perenne* (Figure 3B, see table S5 for ANOVAs summaries). This species contributed to almost 90% of the aboveground mass at the final harvest for all thermo-protocols while we transplanted the same number of individuals as *L. corniculatus*. For the four-species mixture, overyielding was not always due to the overperformance of *L. perenne* only (Figure 3B). Its relative mass decreased from the control thermo-protocol (more than 75% of the mixture aboveground biomass) to the recurrent heatwaves thermo-protocol (less than 40% of the mixture aboveground biomass) while *P*.*lanceolata* had the opposite trend. Under recurrent heatwaves, *P*.*lanceolata* had similar relative aboveground mass in the mixture as *L*.*perenne*. While *L*.*corniculatus* and *F*.*rubra* had low contribution on all thermo-protocols in the four-species mixture, but *L*.*corniculatus* increased slightly its contribution from control conditions to recurrent heatwaves.

### Effects of the mild and severe heatwaves alone or combined on plant responses and leaf parameters

The studied species showed different resistance abilities (during the severe heatwave) and recovery responses (after the severe heatwave) (Figure 4, see Table S6 for t-test summaries). These responses depended on whether they were grown in monocultures or in mixtures and on the event recurrence (Figure 4, see Table S7 for ANOVAs summaries). In terms of resistance, *F. rubra* was resistant only in the four-species mixture during severe HW and recurrent heatwaves; *L. perenne* was more resistant for the severe heatwave on mixtures than monoculture; *L. corniculatus* appeared to be more resistant for the severe heatwave on four-species mixtures than monoculture; *P. lanceolata* had not a clear pattern for the severe heatwave. In terms of recovery, *F. rubra* showed generally recovery in all thermo-protocols in four-species mixture; *L. perenne* showed also generally recovery; *L. corniculatus* showed better recovery on mixtures than monocultures for the severe HWs; *P. lanceolata* showed higher recovery on four-species mixture than monocultures.

**Figure 4.**
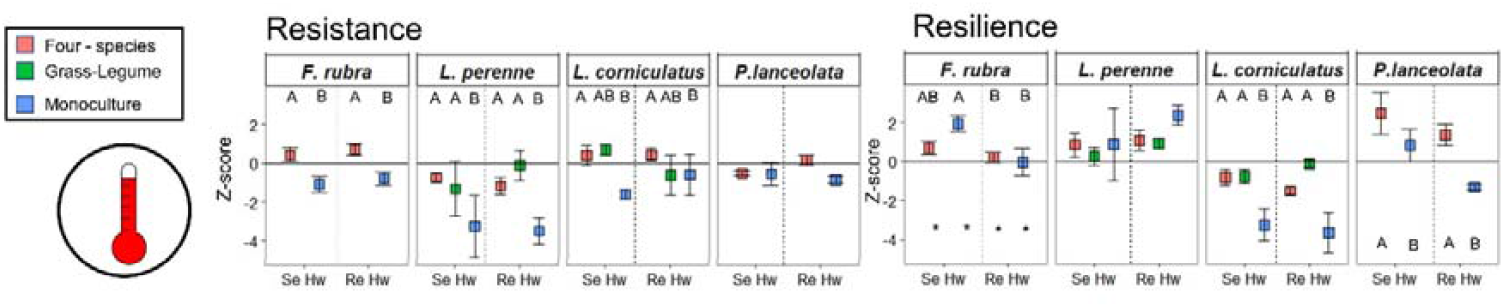
Resistance and recovery to a severe heatwave for each species in monoculture, grass-legume mixture or four-species mixture. Values are Z-scores based on control conditions. Thermo-protocols: **Se**vere heatwave and **Re**current heatwaves. Mean and standard errors. Letters indicate significant differences after multiple comparisons for each species if thermo-protocol and type of cultures were significant on ANOVAs. Asterisks indicate µ≥0 using t-test.

The leaf temperature was the parameter the most impacted by thermo-protocols, culture types and their interactions for all grassland species (Figure 5 and Figure S4, see ANOVA summaries in tables S8). Plants subjected to heatwaves, whatever the thermo-protocol, had higher leaf temperature than control ones. At the first mild heatwaves, at leaf level, *F. rubra* reached an average of 21.03 °C *vs*. 18.99 °C on control plants, *P. lanceolata* 20.52 °C *vs*. 17.57 °C, *L. perenne* 21.21 °C *vs*. 18.72 °C, and *L. corniculatus* 20.96 °C *vs*. 18.19 °C. Notably, *L. perenne* showed higher leaf temperatures on monocultures than in four-species mixture. At the second mild heatwave, at leaf level, *F. rubra* reached an average of 19.58 °C *vs*. 17.22 °C on control plants, *P. lanceolata* 19.95 °C *vs*. 17.60 °C, *L. perenne* 20.30 °C *vs*. 17.72 °C, and *L. corniculatus* 19.43 °C *vs*. 17.26 °C. There were also differences regarding the culture type. *P. lanceolata* showed higher temperature in the four-species mixture than in monocultures. *L. corniculatus* showed lower temperature in monoculture without heatwaves and the higher on grass-legume mixture during heatwaves. At the third heatwave, the severe one, no difference was observed between plants subjected to the severe heatwave alone and to recurrent ones. At leaf level, *F. rubra* reached an average of 32.58 °C *vs*. 16.21 °C on control plants, *P. lanceolata* 32.48 °C *vs*. 16.02°C, *L. perenne* 33.24 °C *vs*. 16.02°C, and *L. corniculatus* 33.4 °C *vs*. 16.06 °C.

**Figure 5.**
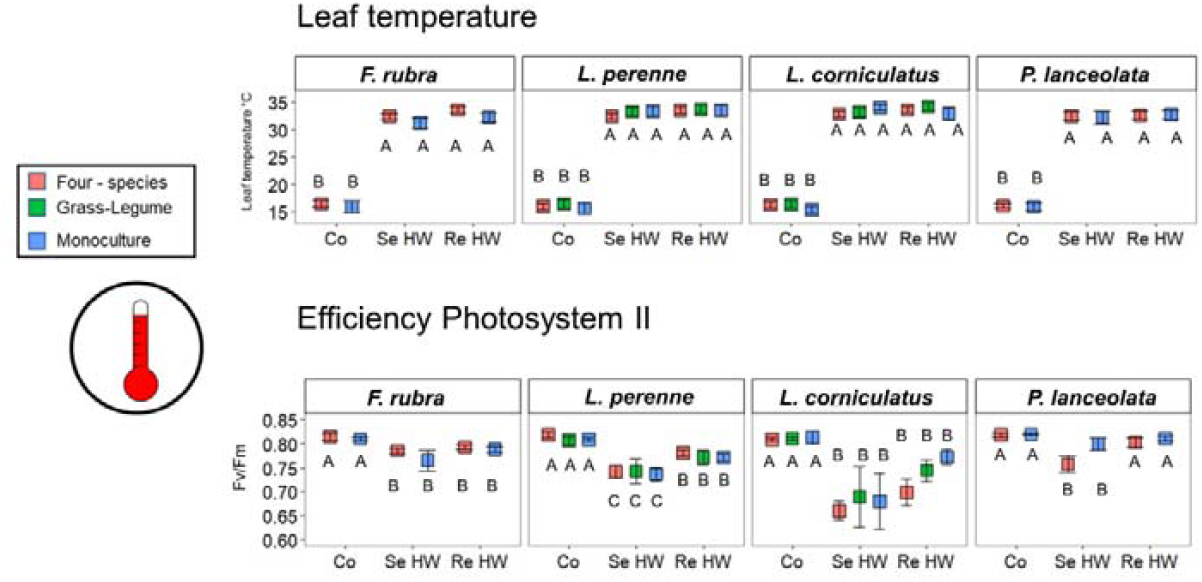
Leaf temperature and efficiency of Photosystem II during a severe heatwave for each species in monoculture, grass-legume mixture or four-species mixture. Thermo-protocols are **Control** chamber (Co), **Se**vere heatwave and **Re**current heatwaves. Mean and standard errors. Capital letters indicate significant differences after multiple comparisons among thermo-protocols, and minor letters among types of cultures.

The efficiency of Photosystem II (Figure 5 and Figure S4), assessed through Fv/Fm index, was only affected during the severe heatwave (see ANOVA summaries in tables S9) for all species and culture types. Nevertheless, *F. rubra* showed worse performance in monocultures during the first mild heatwave (Figure 5), while *P. lanceolata* showed better performance in monocultures than four-species mixture on the heatwave unit. Upon the severe heatwave, there was a significant effect on Fv/Fm, which magnitude differed among species. Nevertheless, this effect was modulated by previous mild heatwaves since the values under recurrent HWs differed from the ones observed under the severe HW alone. *L. perenne* and *P. lanceolata* subjected to mild heatwaves had a higher Fv/Fm than when subjected to only a severe heatwave. *F. rubra, L. perenne, L. corniculatus* had a lower Fv/Fm when subjected to the two thermo-protocols including a severe heatwave (severe HW and recurrent HWs) than control one.

Visual symptoms of heatwave-induced effects were observed just after the severe heatwaves but not after mild heatwaves. These observations indicated that *F. rubra* did not show heatwave-induced visual symptoms, while *P. lanceolata* showed withered leaves, *L. perenne* showed withered leaves and apical senescence, and *L. corniculatus* showed foliar senescence and abscission.

## Discussion

In line with our hypothesis, we demonstrated that the species studied showed different responses to heatwaves. Due to this diversity, in our experiment, when recurrent heatwaves induced overall negative cumulative effects on plant production, mixtures performed better than monocultures. In addition, certain species showed greater resistance or recovery in mixtures than in monocultures, but overall, we did not observe any effect of diversity on leaf parameters. Before delving deeper into these results, it is necessary to understand how the intensity and recurrence of heatwaves affected plant responses.

### The intensity of heatwaves is the main factor to determine the effects

The mild heatwaves, chosen to simulate heatwaves that could occur in spring, had not effect on production, as widely observed in the literature for grassland species in temperate regions (De Boeck *et al*., 2011). In contrast, the severe heatwave, designed to simulate a summer heatwave, was shown to induce negative effects on production for the studied species, as previously observed for grassland species (Wang *et al*., 2008). The difference in induced heat effects between mild heatwaves and severe heatwaves is probably mainly due to the intensity of the heatwave and/or the availability of water used by plants to reduce temperature, as these have been described as the most important factors for heatwaves induced-effects on production (De Boeck *et al*., 2011; Dreesen *et al*., 2014), rather than the timing of occurrence. Nevertheless, the intensity of heatwaves is difficult to compare with other studies due to different conditions for temperature recording (e.g., ambient air temperature, at pot level, at leaf surface…). In our experiment, the set temperatures in the greenhouse used to simulate the thermo-protocols differed notably from the temperature measured at pot level. This substantial temperature difference poses a considerable challenge when trying to decipher the effects of heatwaves. Therefore, the pot level temperature was considered to be the most accurate record of the temperature of the air surrounding plants.

In our scenario (5-day severe heatwave under well-watered conditions), leaf temperatures rose above 32°C, which is the supra-optimal threshold at which *L. perenne* is considered to reduce its leaf expansion rate (Peacock, 1975). Meanwhile, there was no collapse of Photosystem II (PSII), as it did not reach a 50 % drop in activity compared to control conditions. This result can probably be explained by the absence of water stress as the plants were well watered. The strongest inhibition of PSII in *L*.*perenne* was observed when heat events occurred during periods of low soil moisture and not during wet periods (Digrado *et al*., 2017). Furthermore, we observed that leaf temperature (approximately 33°C) was lower than pot-level temperature (approximately 34.5°C), which would indicate high transpiration rates (Blonder & Michaletz, 2018). These high heat-induced transpiration rates relate to the so-called “leaf cooling” phenomenon, which has been widely reported in the literature (Marchin *et al*., 2022). Although the plants studied used this mechanism, we observed only a slight decrease in water content in the system (pot mass loss) during the severe heatwave, indicating that they were all well-watered. This finding is important for understanding the effects of heatwave alone, as it is generally accepted that heat induced damage only occurs in conditions of water shortage (Reynaert *et al*., 2025). In our case, plants managed to maintain lower leaf temperature than air temperature by leaf cooling, but leaf temperature remained above the optimal threshold, resulting in a decline in plant production. This decline can be attributed to the cessation of growth activity rather than to stomatal closure or deterioration of PSII, indicating a dysfunction of carbon sinks rather than carbon sources (Moore *et al*., 2021). Indeed, this suggests that even when there is sufficient water in the system, production will be reduced if heatwaves are more intense than the plant’s threshold for growth activity (Langworthy et al. 2020). Such findings highlight the importance of studying thermal thresholds in the context of the whole source-sink balance, rather than just carbon sources.

### The effect of recurrent events is not the sum of the effect of single events

The lower production observed in the recurrent event thermo-protocol compared to the severe heatwave thermo-protocol suggests a cumulative effect of the individual effects. This cumulative effect has previously been observed in *Poa labillardierei* Steud., a C3 grass native to temperate Australia (French *et al*., 2019), following three heatwave events. A similar cumulative effect was also observed in temperate grassland species when the interval between heatwave events was reduced (Dreesen *et al*., 2014), an interval referred to as the recovery phase, meaning in this case that the longer the phase between events, the better the recovery. To understand the effects of recurrent events, it has been noted that the effects can be species-dependent, but also dependent on the thermal scenario itself, meaning that the intensity, timing, duration and frequency of the event - in relation to the recovery phase - are determining (Bäurle, 2016; Ahrens *et al*., 2021). Overall, in our experiment, all species showed a uniform response, i.e., a negative cumulative effect on production. Our results did not accurately reflect the sum of the individual effects of mild and severe heatwaves, suggesting some form of memory. The mild heatwave induced neutral effects, while the severe heatwave induced a negative effect. Consequently, a previously unobserved negative effect on mild heatwaves became apparent following the occurrence of a severe heatwave. This hidden effect was also observed in another experiment with crop species, where the negative effects of a second event were amplified compared to the first one, maybe due to a lack of time for recovery between events (Angelique *et al*., 2025). Thus, plants may be in a weakened state at the time of the second event (Crisp *et al*., 2016). This is analogous to the phenomenon observed in some temperate grassland species, which suffer leaf drop after the first event and, despite the apparent subsequent recovery, plants show reduced resistance to the following event (Dreesen *et al*., 2014). However, in the present study, only leaf senescence (*L. perenne* and *L. corniculatus*) and even leaf shedding (*L. corniculatus*) were observed during the severe heatwave, but not after the mild heatwaves. It is likely that negative effects not observed during mild heatwaves imprinted uncharacterized physiological or molecular parameters.

Regarding the efficiency of Photosystem II, we observed a priming effect in *L. perenne* and *P. lanceolata*, as Fv/Fm ratio were higher on plants exposed to previous mild heatwaves during the severe heatwave. This priming effect leads to an acclimation of Fv/Fm ratio, which has already been evidenced in different scenarios of recurrent heatwave events (Zhu *et al*., 2018; French *et al*., 2019). Hence, the efficiency of Photosystem II can adapt to heatwaves (Geange *et al*., 2021). These results, both a cumulative negative effect on production and an acclimation of the Photosystem II, again lines on the importance to evaluate the whole carbon sink-source balance in the face of recurrent heatwaves.

### Buffering negative effects by multi-species assemblages

The attenuation of the effects of drought events by plant diversity is well-known, even in intensively managed temperate grasslands (Hofer *et al*., 2016; Grange *et al*., 2021; Haughey *et al*., 2023). However, before our study, no evidence showed that multi-species assemblages could buffer heatwave effects. While grassland community responses to heatwaves had been studied (De Boeck *et al*., 2011; Dreesen *et al*., 2012, 2014), comparison with monocultures were lacking. We observed overyielding, though it depended on mixture type and, in four-species mixtures, on thermo-protocols. Overyielding under climatic events may result from synergistic species interactions (Haughey *et al*., 2023), the complementarity (Barry *et al*., 2019), or overperformance of dominant species (Hofer *et al*., 2016), the sampling effect (Loreau, 1998). The four-species, grass-legume mixtures and *L*.*perenne* monoculture showed the highest yields under all thermo-protocols, mainly due to *L. perenne*. Under recurrent heatwaves, though *P*.*lanceolata* monocultures showed lower yield, the higher four-species mixture yield was not only due to *L. perenne* but also to the high contribution of *P. lanceolata*, especially under recurrent heatwaves. In addition, *L. corniculatus* had a higher contribution under recurrent heatwaves thermo-protocol. Both species had better recovery to recurrent heatwave in the four-species mixture than in the monocultures. Overyielding under recurrent heatwaves may be achieved through abiotic facilitation (Barry *et al*., 2019), following the poorer performance of *L. perenne*. This abiotic facilitation may be achieved by some plants providing resources to neighbors, or by buffering induced-event effects (Barry *et al*., 2019). The interaction between *P*.*lanceolata* and N-fixing species is known (Temperton *et al*., 2007). It may be that the poorer performance of *L. perenne* allows *P. lanceolata* to use a resource usually monopolized by *L. perenne* and offered by *L. corniculatus*. From the other group of abiotic facilitation mentioned, processes that promote microclimate buffering have been suggested in the context of heat events (Reynaert *et al*., 2025). In our study, we measured both leaf temperature and Fv/Fm index to observe whether microclimatic buffering occurred. In general, neither leaf temperature nor Fv/Fm values of the studied species showed a better performance in mixtures than in monocultures during severe heatwaves. However, *F*.*rubra, L*.*perenne* and *L*.*corniculatus* showed better resistance in mixtures than in monocultures, suggesting microclimatic buffering processes between species, that remain to be identified. One perspective could be to measure leaf temperature not only in the upper third of plants, as we did, because changes in branch angle or canopy openness can affect temperature (Blonder & Michaletz, 2018). The lower third or the basal part, even belowground, are important for the growth activity in grassland species (Klimešová *et al*., 2016) and could be relevant points for temperature measurements.

## Acknowledgements

We are grateful to Barenbrug and Ceriance, both companies provided the seeds used in this experiment. Anne-Françoise Ameline, Véronique Signoret, Magali Bodereau and Josiane Pichon for their work in greenhouse management and samples processing. Jean-Bernard Cliquet, Jean-François Odoux, Leonidas Kougiteas, Juliette Senecal, Mona Brancadoro, Enora Chauvin and Paul Laporte for help with mowing and samples processing. Til Feike to provide useful comments on the experimental design.

## Author contributions

All authors designed and set up the experiment; AC, maintained the experiment, measured all variables, analysed data and led the manuscript writing; All authors discussed the results and wrote the manuscript

## Data availability

The data that support the findings of this study are available from the corresponding author, AC, upon reasonable request.

