## Supplementary files for "Temperate grasslands facing heatwaves: species diversity buffers effects on shoot growth but not on leaf parameters"

Table S1. ANOVA summaries for Pot mass variation on each heatwave. Type unit has two levels: control unit and heatwave (HW) unit. Type Thermo-protocol has two levels: with or without Mild HWs. We use type II errors to evaluate significant differences.

| | Types | Chisq | df | p-value |
|---|---|---|---|---|
| Mild HW1 | Unit | 0.31 | 1 | 0.578 |
| Mild HW 2 | Unit | 3.06 | 1 | 0.080 |
| Severe HW | Unit | 71.59 | 1 | 0.000 |
| | Thermo-protocol | 0.97 | 1 | 0.324 |

Table S2. ANOVA summaries of the effect on mass accumulation. We use type II errors to evaluate significant differences.

| | Types | Chisq | Df | P value |
|---|---|---|---|---|
| Sum of mowing | Mild HW | 2.26 | 1 | 0.133 |
| | Severe HW | 1.91 | 1 | 0.167 |
| | Culture | 406.09 | 5 | 0.000 |
| | Mild HW : Severe HW | 37.71 | 1 | 0.000 |
| | Severe HW : Culture | 5.67 | 5 | 0.340 |
| | Mild HW : Culture | 7.61 | 5 | 0.179 |
| | Mild : Severe : Culture | 15.11 | 5 | 0.010 |
| 1st mowing | Culture | 198.61 | 5 | 0.000 |
| 2nd mowing | Heatwave | 1.51 | 1 | 0.220 |
| | Culture | 185.10 | 5 | 0.000 |
| | Mass at 1st mowing | 33.35 | 1 | 0.000 |
| | HW : Culture | 2.52 | 5 | 0.772 |
| 3rd mowing | Mild HW | 6.02 | 1 | 0.014 |
| | Severe HW | 78.69 | 1 | 0.000 |
| | Culture | 87.07 | 5 | 0.000 |
| | Mass at 1st mowing | 1.48 | 1 | 0.224 |
| | Mass at 2nd mowing | 46.42 | 1 | 0.000 |
| | Mild HW : Severe HW | 14.80 | 1 | 0.000 |
| | Mild HW : Culture | 5.55 | 5 | 0.353 |
| | Severe HW : Culture | 11.01 | 5 | 0.051 |
| | Mild : Severe : Culture | 4.40 | 5 | 0.494 |

Table S3. Estimated means different to 0 on non-transgressive overyielding.

| | | Emmean | p-value |
|---|---|---|---|
| Sum of mowing | | | |
| Grass-Legume | | | |
| | Control | 0.274 | 0.001 |
| | Mild HWs | 0.314 | 0.001 |
| | Severe HW | 0.102 | 0.052 |
| | Recurrent HWs | 0.312 | 0.001 |
| Four species | | | |
| | Control | 0.254 | 0.001 |
| | Mild HWs | 0.313 | 0.001 |
| | Severe HW | 0.126 | 0.006 |
| | Recurrent HWs | 0.272 | 0.001 |
| 1st mowing | | | |
| Grass-Legume | | | |
| | Control | 0.095 | 0.011 |
| Four species | | | |
| | Control | 0.204 | 0.001 |
| 2nd mowing | | | |
| Grass-Legume | | | |
| | Control | 0.297 | 0.001 |
| | Mild HWs | 0.409 | 0.001 |
| Four species | | | |
| | Control | 0.174 | 0.001 |
| | Mild HWs | 0.220 | 0.001 |
| 3rd mowing | | | |
| Grass-Legume | | | |
| | Control | 0.367 | 0.001 |
| | Mild HWs | 0.341 | 0.001 |
| | Severe HW | 0.314 | 0.001 |
| | Recurrent HWs | 0.427 | 0.001 |
| Four species | | | |
| | Control | 0.357 | 0.001 |
| | Mild HWs | 0.339 | 0.001 |
| | Severe HW | 0.257 | 0.001 |
| | Recurrent HWs | 0.378 | 0.001 |

Table S4. Estimated means different to 0 on transgressive overyielding.

| Sum of mowing | | Emmean | p-value |
| --- | --- | --- | --- |
| Grass-Legume | | | |
| | Control | 0.027 | 0.515 |
| | Mild HWs | 0.103 | 0.022 |
| | Severe HW | -0.053 | 0.215 |
| | Recurrent HWs | 0.101 | 0.025 |
| Four species | | | |
| | Control | -0.056 | 0.101 |
| | Mild HWs | 0.083 | 0.018 |
| | Severe HW | -0.056 | 0.104 |
| | Recurrent HWs | -0.034 | 0.316 |
| 1st mowing | | | |
| Grass-Legume | | | |
| | Control | 0.044 | 0.139 |
| Four species | | | |
| | Control | -0.037 | 0.147 |
| 2nd mowing | | | |
| Grass-Legume | | | |
| | Control | -0.024 | 0.532 |
| | Mild HWs | 0.097 | 0.020 |
| Four species | | | |
| | Control | -0.131 | 0.001 |
| | Mild HWs | -0.069 | 0.052 |
| 3rd mowing | | | |
| Grass-Legume | | | |
| | Control | -0.010 | 0.817 |
| | Mild HWs | 0.065 | 0.087 |
| | Severe HW | -0.067 | 0.081 |
| | Recurrent HWs | -0.038 | 0.311 |
| Four species | | | |
| | Control | -0.049 | 0.268 |
| | Mild HWs | 0.031 | 0.491 |
| | Severe HW | -0.036 | 0.415 |
| | Recurrent HWs | -0.155 | 0.001 |

Table S5a. ANOVAs summaries of the effect on relative mass accumulation among species per each thermo-protocol and type of mixture.

| | ~species | Control | Mild heatwaves | Severe heatwave | Recurrent heatwave |
|---|---|---|---|---|---|
| Grass-Legume | *Chisq* | 1712.1 | 61.06 | 1655.5 | 63.55 |
| | p-value | 0.000 | 0.000 | 0.000 | 0.000 |
| Four-species | *Chisq* | 481.1 | 55.95 | 98.35 | 46.38 |
| | p-value | 0.000 | 0.000 | 0.000 | 0.000 |

Table S5b. ANOVAs summaries of the effect on relative mass accumulation among thermo-protocols per each species and type of mixture.

| | ~thermoprotocol | *F.rubra* | *L.perenne* | *L.corniculatus* | *P.lanceolata* |
|---|---|---|---|---|---|
| Grass-Legume | Chisq | | 3.69 | 3.69 | |
| | p-value | | 0.297 | 0.297 | |
| Four-species | Chisq | 3.83 | 12.16 | 10.70 | 14.32 |
| | p-value | 0.281 | 0.007 | 0.013 | 0.002 |

Table S6. Estimate means different to 0 of the effect on resistance and recovery responses per each event, thermo-protocol, species and type of culture.

| Event | Thermo | Species | Culture | Resistance Estimate | p-value | Recovery Estimate | p-value |
|---|---|---|---|---|---|---|---|
| Mild heatwaves | | *F.rubra* | Four-species | 0.462 | 0.229 | 0.668 | 0.040 |
| | | | Monoculture | -1.671 | 0.005 | -0.393 | 0.082 |
| | | *L.perenne* | Four-species | 0.067 | 0.959 | -1.550 | 0.040 |
| | | | Grass-Legume | 2.556 | 0.095 | -2.493 | 0.006 |
| | | | Monoculture | 0.988 | 0.591 | -0.155 | 0.881 |
| | | *L.corniculatus* | Four-species | 0.858 | 0.012 | -0.405 | 0.113 |
| | | | Grass-Legume | 1.237 | 0.002 | -0.493 | 0.096 |
| | | | Monoculture | 0.549 | 0.238 | -1.222 | 0.002 |
| | | *P.lanceolata* | Four-species | -0.080 | 0.728 | -0.628 | 0.153 |
| | | | Monoculture | -0.268 | 0.412 | -0.317 | 0.602 |
| Severe heatwave | Severe | *F.rubra* | Four-species | 0.427 | 0.210 | 0.705 | 0.040 |
| | | | Monoculture | -1.108 | 0.035 | 1.950 | 0.001 |
| | | *L.perenne* | Four-species | -0.733 | 0.373 | 0.854 | 0.236 |
| | | | Grass-Legume | -1.316 | 0.174 | 0.262 | 0.747 |
| | | | Monoculture | -3.254 | 0.012 | 0.882 | 0.381 |
| | | *L.corniculatus* | Four-species | 0.397 | 0.273 | -0.813 | 0.052 |
| | | | Grass-Legume | 0.674 | 0.115 | -0.752 | 0.111 |
| | | | Monoculture | -1.618 | 0.005 | -3.262 | 0.001 |
| | | *P.lanceolata* | Four-species | -0.526 | 0.048 | 2.48 | 0.019 |
| | | | Monoculture | -0.559 | 0.119 | 0.84 | 0.514 |
| | Recurrent | *F.rubra* | Four-species | 0.699 | 0.022 | 0.213 | 0.518 |
| | | | Monoculture | -0.787 | 0.055 | -0.022 | 0.961 |
| | | *L.perenne* | Four-species | -1.173 | 0.028 | 1.086 | 0.013 |
| | | | Grass-Legume | -0.109 | 0.847 | 0.938 | 0.052 |
| | | | Monoculture | -3.501 | 0.001 | 2.388 | 0.001 |
| | | *L.corniculatus* | Four-species | 0.428 | 0.488 | -1.502 | 0.001 |
| | | | Grass-Legume | -0.633 | 0.376 | -0.124 | 0.754 |
| | | | Monoculture | -0.587 | 0.500 | -3.664 | 0.001 |
| | | *P.lanceolata* | Four-species | 0.143 | 0.511 | 1.38 | 0.010 |
| | | | Monoculture | -0.853 | 0.018 | -1.30 | 0.058 |

Table S7a. ANOVAs summaries of the effect on resistance among type of culture per each species and type of culture

| | Mild heatwaves | | Severe heatwave | | | | | |
|---|---|---|---|---|---|---|---|---|
| | ~culture | | culture | | thermo | | cult*thermo | |
| | Chisq | p-value | Chisq | p-value | Chisq | p-value | Chisq | p-value |
| F.rubra | 10.92 | 0.001 | 18.61 | 0.000 | 0.76 | 0.383 | 0.01 | 0.944 |
| L.perenne | 1.61 | 0.448 | 11.63 | 0.003 | 0.06 | 0.806 | 1.47 | 0.480 |
| L.corniculatus | 1.42 | 0.492 | 6.42 | 0.040 | 0.17 | 0.676 | 3.60 | 0.165 |
| P.lanceolata | 0.23 | 0.631 | 3.66 | 0.056 | 1.89 | 0.169 | 3.20 | 0.074 |

Table S7b. ANOVAs summaries of the effect on resilience among type of culture per each species and type of culture

| | Mild heatwaves | | Severe heatwave | | | | | |
|---|---|---|---|---|---|---|---|---|
| | ~culture | | culture | | thermo | | cult*thermo | |
| | Chisq | p-value | Chisq | p-value | Chisq | p-value | Chisq | p-value |
| F.rubra | 8.10 | 0.004 | 1.82 | 0.177 | 7.80 | 0.005 | 3.92 | 0.048 |
| L.perenne | 3.12 | 0.210 | 2.08 | 0.354 | 1.59 | 0.207 | 0.87 | 0.65 |
| L.corniculatus | 3.88 | 0.144 | 4.44 | 0.000 | 0.30 | 0.584 | 2.99 | 0.22 |
| P.lanceolata | 0.18 | 0.672 | 6.61 | 0.010 | 3.35 | 0.067 | 0.38 | 0.536 |

Table S8. ANOVA summaries for leaf temperature per each species, type of culture and thermo-protocol.

| | | 1st Mild heatwave | | 2nd Mild heatwave | | Severe heatwave | |
|---|---|---|---|---|---|---|---|
| | Type | Chisq | p-value | Chisq | p-value | Chisq | p-value |
| *F. rubra* | Heatwave | 26.33 | 0.000 | 43.15 | 0.000 | 948.8 | 0.000 |
| | Culture | 3.79 | 0.051 | 0.01 | 0.926 | 3.72 | 0.054 |
| | Heatwave:culture | 1.23 | 0.267 | 0.31 | 0.579 | 0.575 | 0.750 |
| *P. lanceolata* | Heatwave | 96.90 | 0.000 | 57.57 | 0.000 | 709.0 | 0.000 |
| | Culture | 1.37 | 0.243 | 8.13 | 0.004 | 0.139 | 0.709 |
| | Heatwave:culture | 0.15 | 0.697 | 6.95 | 0.008 | 0.192 | 0.909 |
| *L. corniculatus* | Heatwave | 57.06 | 0.000 | 63.88 | 0.000 | 1684.5 | 0.000 |
| | Culture | 1.02 | 0.601 | 12.47 | 0.002 | 1.32 | 0.516 |
| | Heatwave:culture | 0.45 | 0.800 | 4.80 | 0.091 | 2.00 | 0.735 |
| *L. perenne* | Heatwave | 290.59 | 0.000 | 81.55 | 0.000 | 1439.6 | 0.000 |
| | Culture | 17.83 | 0.000 | 0.31 | 0.856 | 1.04 | 0.596 |
| | Heatwave:culture | 1.54 | 0.463 | 2.46 | 0.293 | 1.35 | 0.853 |

Table S9. ANOVA summaries for Fv/Fm per each species, type of culture and thermo-protocol.

| | | 1st Mild heatwave | | 2nd Mild heatwave | | Severe heatwave | |
|---|---|---|---|---|---|---|---|
| | Type | Chisq | p-value | Chisq | p-value | Chisq | p-value |
| *F. rubra* | Heatwave | 0.83 | 0.363 | 0.14 | 0.706 | 21.30 | 0.000 |
| | Culture | 12.38 | 0.000 | 1.84 | 0.175 | 2.24 | 0.135 |
| | Heatwave:culture | 4.77 | 0.029 | 0.28 | 0.598 | 1.20 | 0.550 |
| *P. lanceolata* | Heatwave | 1.76 | 0.185 | 4.85 | 0.028 | 26.26 | 0.000 |
| | Culture | 0.57 | 0.449 | 4.85 | 0.028 | 1.71 | 0.192 |
| | Heatwave:culture | 0.90 | 0.344 | 0.61 | 0.436 | 2.01 | 0.367 |
| *L. corniculatus* | Heatwave | 0.44 | 0.506 | 2.10 | 0.147 | 31.39 | 0.000 |
| | Culture | 1.05 | 0.590 | 4.93 | 0.086 | 3.30 | 0.192 |
| | Heatwave:culture | 4.74 | 0.093 | 1.12 | 0.571 | 1.63 | 0.804 |
| *L. perenne* | Heatwave | 0.13 | 0.721 | 0.22 | 0.640 | 55.96 | 0.000 |
| | Culture | 5.32 | 0.070 | 0.37 | 0.832 | 0.89 | 0.639 |
| | Heatwave:culture | 11.02 | 0.004 | 11.18 | 0.004 | 0.37 | 0.985 |

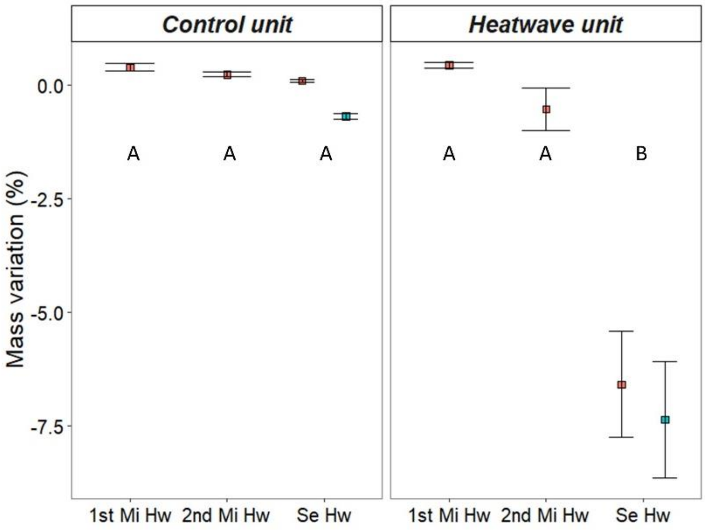


Figure S1. Pot mass loss. Mean and standard errors. Blue squares represent pots previously subjected to mild heatwaves and red pots only subjected to severe heatwave. Capital letters indicate significant differences after multiple comparisons.

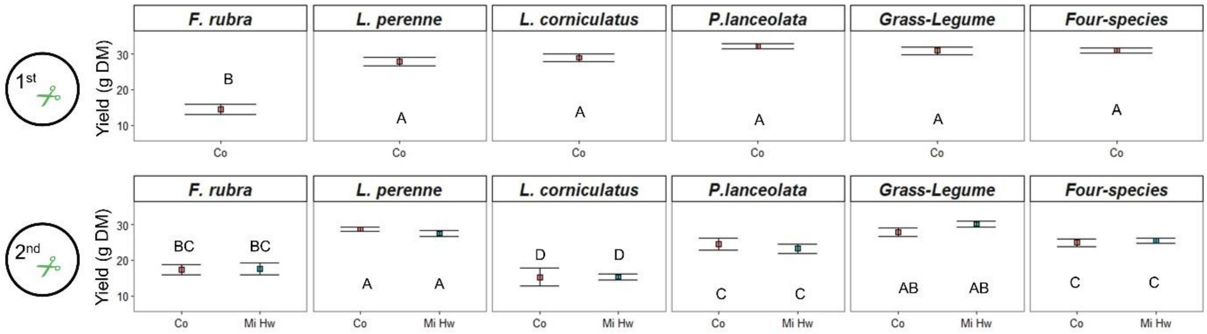


Figure S2. Mass production (g Dry Mass) for 1st and 2nd mowing, culture type (monocultures, grass-legume and four species mixtures) and thermo-protocol (Co: control, Mi Hw: mild heatwaves, Se Hw: severe heatwave, Re Hw: recurrent heatwaves). Means and standard error are shown. Capital letters indicate significant difference among culture types after multiple comparisons, and minor letters indicate difference among thermo-protocols for each culture type ($p<0.05$).

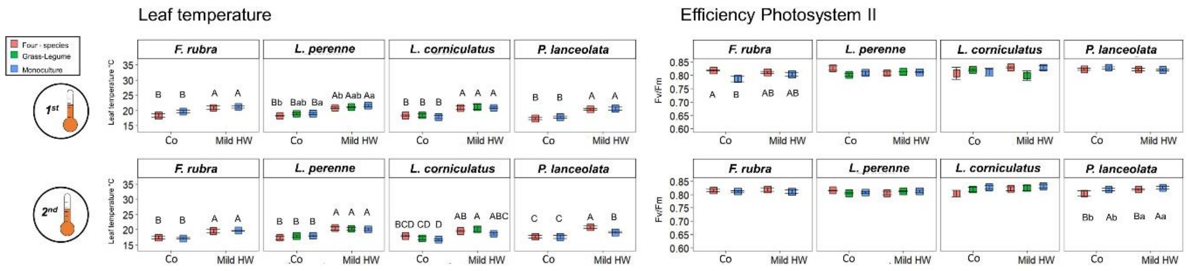


Figure S3. Leaf temperature and efficiency of Photosystem II during mild heatwaves for each species in monoculture, grass-legume mixture or four-species mixture. Heatwave-treatments are **Control** chamber (Co), **Se**vere heatwave and **Re**current heatwaves. Mean and standard errors. Capital letters indicate significant differences after multiple comparisons among thermo-protocols, and minor letters among types of cultures.
